# A comparison of single and multi-echo processing of fMRI data during overt autobiographical recall

**DOI:** 10.1101/2021.11.03.467128

**Authors:** Adrian W. Gilmore, Anna M. Agron, Estefanía I. González-Araya, Stephen J. Gotts, Alex Martin

## Abstract

Recent years have seen an increase in the use of multi-echo fMRI designs by cognitive neuroscientists. Acquiring multiple echoes allows one to reduce thermal noise and identify nuisance signal components in BOLD data (Kundu et al., 2012). At the same time, multi-echo acquisitions increase data processing complexity and may incur a cost to the temporal and spatial resolution of the acquired data. Here, we re-examine a multi-echo dataset (Gilmore et al., 2021a) analyzed using multi-echo ICA (ME-ICA) and focused on hippocampal activity during the overly spoken recall of recent and remote autobiographical memories. The goal of the present series of analyses was to determine if ME-ICA’s theoretical denoising benefits might lead to a practical difference in the overall conclusions reached. Compared to single echo data, ME-ICA led to qualitatively different conclusions regarding hippocampal contributions to autobiographical recall: whereas the single echo analysis largely failed to reveal hippocampal activity relative to an active baseline, ME-ICA results supported predictions of the Standard Model of Consolidation and a time limited hippocampal involvement (Alvarez and Squire, 1994). These data provide a practical example of the benefits multi-echo denoising in a naturalistic memory paradigm and demonstrate how they can be used to address long-standing theoretical questions.

## BODY TEXT

Naturalistic fMRI paradigms seek to improve our understanding of the neural bases of “everyday” behavior and strive to be less artificial than more traditional laboratory paradigms (e.g., Hasson and Honey, 2012; Haxby et al., 2020). Naturalistic experiments might involve scanning participants while they watch or describe a popular television show (Chen et al., 2017), read complex narrative passages in the scanner (Finn et al., 2018), or engage in spontaneous conversation with another individual (Jasmin et al., 2019). Although naturalistic paradigms offer opportunities to study brain-behavior relationships beyond those observable in more tightly controlled paradigms, they can also pose additional challenges.

For studies involving spoken responses, the basic act of speaking in an fMRI environment represents a potential issue. Speech will necessarily produce head motion, and if speech is continuous (e.g., Chen et al., 2017; Jasmin et al., 2019) then approaches that require responses between volume acquisitions (e.g., Gracco et al., 2005) or censor specific frames containing speech data (Siegel et al., 2013) are not applicable. Instead, one might be better served by applying recent fMRI timeseries denoising approaches, such as multi-echo ICA (ME-ICA), to remove nuisance signal from one’s data (see Caballero-Gaudes and Reynolds, 2017).

ME-ICA involves decomposing the multi-echo timeseries into ICA components that are identified as “BOLD-like” and “noise-like”, combining the signal across multiple echoes in each TR, and subsequently regressing the noise-like timeseries identified in ICA from the combined data (for in-depth treatments, see Kundu et al., 2012; Kundu et al., 2013; Gonzalez-Castillo et al., 2016; Kundu et al., 2017; Power et al., 2018). Although ME-ICA is employed to improve overall data quality, its practical benefit should be balanced against its costs. In this case, the tradeoffs may consist of slower TRs, reduced coverage, and/or larger voxel resolutions to allow for the additional readouts within each functional volume. In addition, techniques such as ME-ICA require additional preprocessing steps that may not be implemented in all analysis packages.

In this report, the practical benefits of ME-ICA processing are assessed using a recently acquired naturalistic dataset intended to study human memory function (Gilmore et al., 2021a). Forty participants freely and overtly recalled recent and remote autobiographical events for periods of approximately two minutes while undergoing fMRI. As a control task, participants were asked to verbally describe complex photographs. One notable finding from these data was evidence supporting a temporally graded and time-limited role of the hippocampus in the recall of autobiographical memories—an issue that has been discussed at length with several established “camps” in the literature (for recent reviews of various hypotheses, see Squire et al., 2015; Barry and Maguire, 2019; Yonelinas et al., 2019; Gilboa and Moscovitch, 2021). Overt recall was employed to provide experimental knowledge of the type of information being retrieved during recall (Gilmore et al., 2021b) as this, along with the age of a recalled memory, appears to be a critical variable in the debate of hippocampal contributions to remote recall. In the present report, results of the ME-ICA processed data were compared to a more standard processing stream that utilizes a single echo (SE), matched for typical scan acquisition parameters that might be used in any number of memory studies (Table 1; for additional information on the processing and modeling of fMRI data, see Detailed Methods). If the SE results match those of the ME-ICA processed data, it might suggest that multi-echo acquisitions, despite their benefits in some situations, may not be worth their inherent costs when applied to datasets involving continuous speech. On the other hand, if the ME-ICA processed data provided results divergent from an SE approach, then the results would provide a clear example of the benefits of an advanced denoising strategy employed in a naturalistic paradigm.

**Table 1.** Summary of analysis pipelines compared in this report.

| **Standard Processing (2<sup>nd</sup> echo only)** | **ME-ICA denoising (3 echoes)** |
| --- | --- |
| First 4 frames removed | First 4 frames removed for each TE |
| Timeseries despiked | Timeseries despiked for each TE |
| Slice-time corrected | Slice-time corrected for each TE |
| Volume registration (rigid body) | Volume registration for each TE |
|  | ICA denoising: identifies “BOLD-like” and “noise-like” components across TEs |
|  | “Optimally combined” linear combination of TEs weighted by each voxel’s T2* |
|  | “Noise-like” components regressed from the optimally combined data |
| EPI registered to MP-RAGE | Optimally combined data registered to MP-RAGE |
| Timeseries data smoothed and converted to % signal change | Timeseries data smoothed and converted to % signal change |
| Data registered to Talairach atlas (TT N27) | Data registered to Talairach atlas (TT N27) |
| GLM-based mass univariate analysis | GLM-based mass univariate analysis |

Within the hippocampus, anterior and posterior hippocampal subregions (Figure 1A) were manually defined for each participant as regions of interest (ROIs) and activity in each condition was averaged for all voxels within each ROI (Figure 1B, C). A repeated measures ANOVA, with factors of temporal distance (2 levels: Today, 5-10 years ago), subregion (2 levels: anterior, posterior), hemisphere (2 levels: left, right), and pipeline (2 levels: SE, ME-ICA) was employed. There was a significant main effect of pipeline, *F*(1,39) = 6.83, *p* = .013, reflecting the larger BOLD response magnitudes for the SE processed data. This is consistent with prior investigations comparing SE and ME-ICA processing, which have reported attenuated BOLD signal change estimates despite the overall improvement in contrast-to-noise ratios (Gonzalez-Castillo et al., 2016). No other significant main effects were observed (*p*s ≥ .276). Consistent with a prior report using these data (Gilmore et al., 2021a), there was a single 2-way interaction of subregion and temporal distance, *F*(1,39) = 10.87, *p* = .002, reflecting different patterns of activity observed in anterior and posterior hippocampal subregions as a function of event recency (other 2-way interaction *p*s ≥ .138). However, and critically, this interaction must be qualified by a 3-way interaction among the factors of pipeline, subregion, and temporal distance, *F*(2,78) = 4.48, *p* = .041, (other 3-way interaction *p*s ≥ .305). Unpacking this result revealed that significantly greater activity for recent (Today) than remote (5-10 year ago) activity was present in posterior, and not anterior, hippocampal regions, but only for the ME-ICA processed data (anterior hippocampus: *t*(39) = .717, *p* = .478; posterior hippocampus: *t*(39) = 2.91, *p* = .006) and not the SE data (anterior hippocampus: *t*(39) = .876, *p* = .386; posterior hippocampus: *t*(39) = 1.43, *p* = .160). That is, a finding of temporally graded activity relied upon the improved denoising afforded by ME-ICA. No 4-way interaction was observed, *F*(1,39) = .05, *p* = .824).

**Figure 1.**
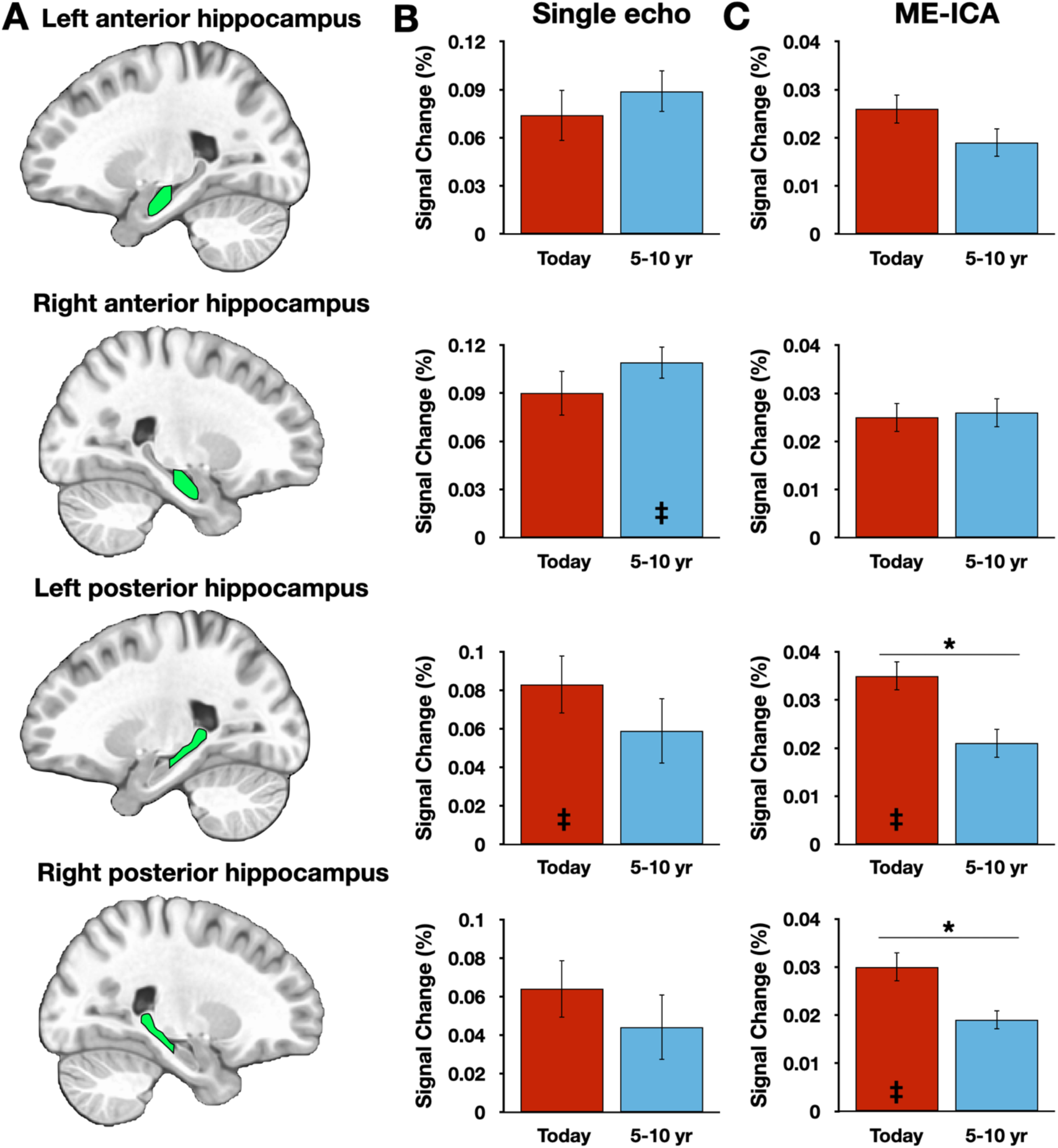
Hippocampal results vary across preprocessing streams. A) Graphic depictions of each manually defined hippocampal subregion. B) Magnitude estimates from the single echo analysis pipeline, plotted against the Picture Description control task. C) Magnitude estimates from the multi-echo ICA pipeline, plotted against the Picture Description control task. Inset double daggers signify a significant difference from the Picture Description control task (FDR-corrected for multiple comparisons). Error bars reflect within-subject standard error.

Hippocampal response magnitudes were then compared to the Picture Description baseline task, as done previously. A time-limited role, as suggested by the Standard Model of Consolidation (Squire and Alvarez, 1995; Squire et al., 2015) would predict significant activations for recent, but not remote, time periods, whereas hypotheses that predict a continuous involvement (e.g., Nadel and Moscovitch, 1997; Sekeres et al., 2018) would predict consistent engagement over the baseline ask. Only 4 of the 16 activations were significant following Bonferroni correction (achieving *p* < .0031): 2 for ME-ICA datapoints associated with the Today condition (Right posterior hippocampus, *t*(39) = 3.90, *p* = 0 .0004; left posterior hippocampus, *t*(39) = 3.38, p = .0016) and two SE datapoints, one for the 5-10 year ago condition (Right anterior hippocampus, *t*(39) = 3.61, p = .0009) and one for the Today condition (left posterior hippocampus, *t*(39) = 3.34, p = .0019) (Figure 1B,C). Applying an FDR correction (Benjamini and Yekutieli, 2001) instead of Bonferroni did not alter this pattern of results.

Basic conclusions regarding hippocampal participation in recent and remote recall therefore differ between SE and ME-ICA processing streams—whereas the ME-ICA data reflected a temporally graded and time-limited hippocampal role during recall, no clear support for any model was revealed by the SE analysis. However, beneficial effects of ME-ICA processing would be expected at the whole-brain level as well. Thus, a voxelwise contrast of activity related to the Today and 5-10 year ago conditions (paired-samples, two-tailed) was performed separately on the SE and ME-ICA data. The SE analysis identified a large cluster in medial parietal cortex, with local maxima in the mid/posterior cingulate cortex and bilaterally in the precuneus, as well as bilateral inferior parietal lobule clusters (Figure 2A; Table 2). Largely convergent results were obtained following ME-ICA processing, although commonly-identified clusters were larger and additional significant clusters were identified in left frontal cortex and right lateral temporal cortex (Figure 2B,C; Table 2). No identified clusters were unique to the SE data.

**Table 2.**
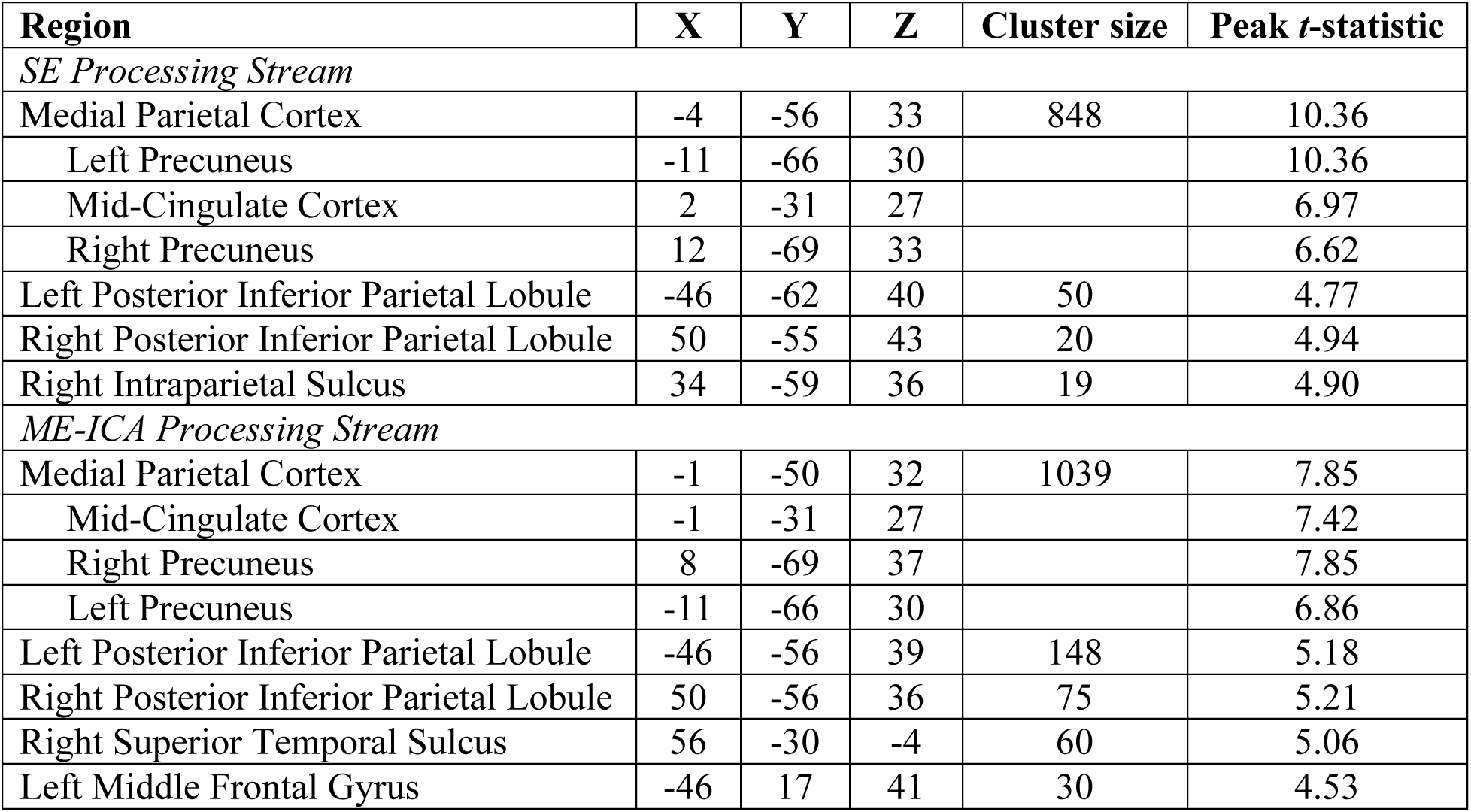
Regions identified in the voxelwise analysis of temporal distance effects for SE and ME-ICA processing streams. Medial parietal subregions reflect discrete local maxima within a larger cluster. Coordinates refer to centers of mass in MNI152 space.

**Figure 2.**
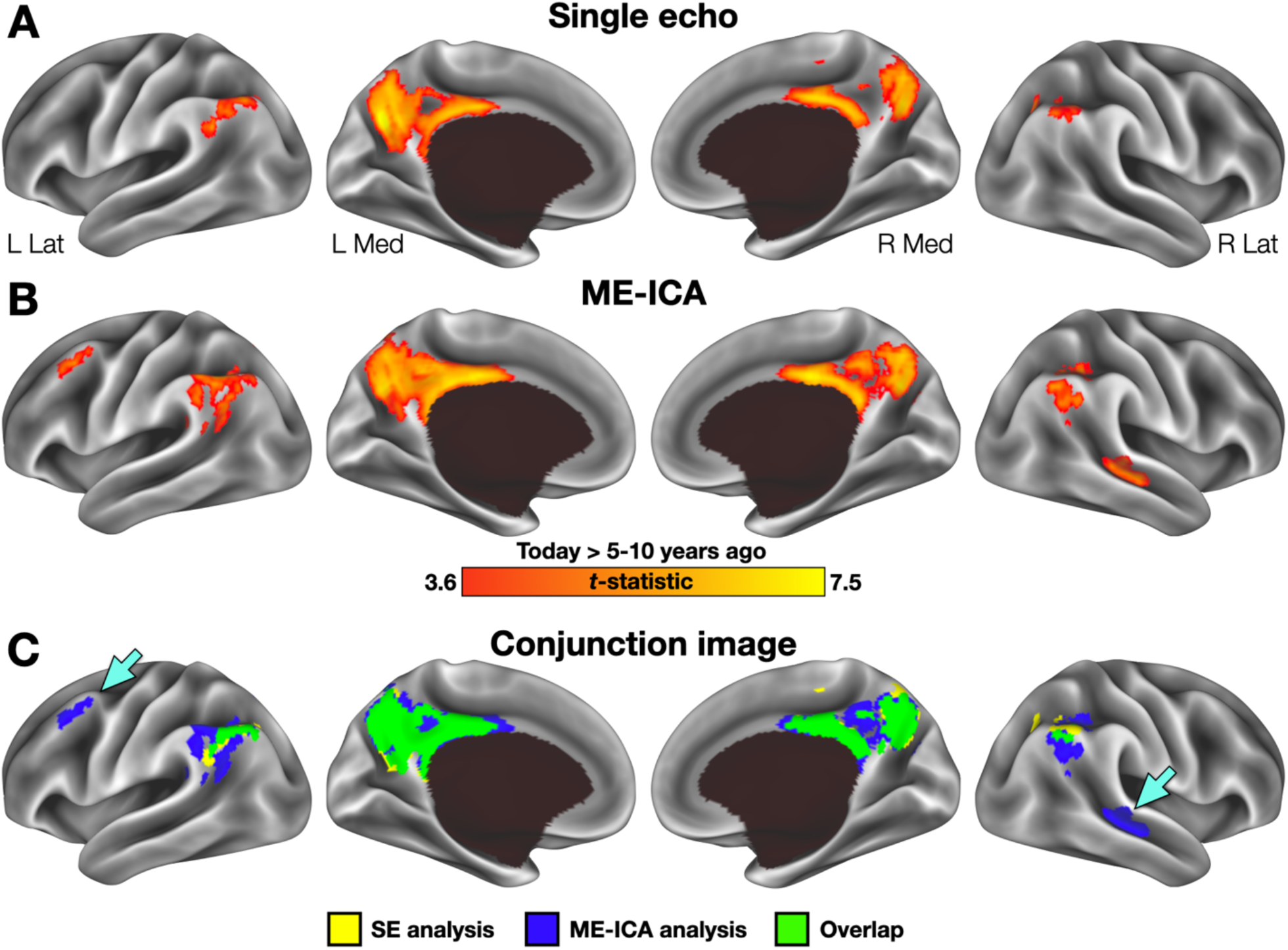
Voxelwise results across preprocessing streams. A) Regions exhibiting greater activity in the Today than 5-10 year ago condition following SE preprocessing included a large cluster in the posterior midline and inferior parietal lobule bilaterally. No clusters were identified with significantly greater activity for the 5-10 year ago condition. B) Regions identified in the same analysis using ME-ICA data were larger and included several additional clusters in the right temporal and left frontal cortex. C) A binarized conjunction image allows for easy visualization of the improved sensitivity offered by ME-ICA processing. Inset arrows identify significant clusters absent from the SE analysis.

In this report, we sought to investigate the practical benefits of an advanced fMRI denoising technique under naturalistic recall conditions. Given the rise in the number of experiments seeking to implement more naturalistic conditions, comparing SE and ME-ICA approaches to data processing might help investigators optimize their sequence and processing pipeline selections. At least in the context of the current dataset, the results of ME-ICA denoising were encouraging.

The general efficacy of ME-ICA in denoising fMRI data has been documented elsewhere (Gonzalez-Castillo et al., 2016), particularly in investigations of functional connectivity (Kundu et al., 2012; Kundu et al., 2013; Kundu et al., 2017; Power et al., 2018; see also Spreng et al., 2019; Power et al., 2019). The purpose of this investigation was not to retread this same ground, but instead to focus on the practical utility of ME-ICA denoising in a naturalistic recall paradigm—an approach in which sophisticated denoising may be particularly important. In this case, ME-ICA-derived improvements appear to have been necessary for the findings obtained previously. This is most clearly demonstrated in the hippocampal results. Data processed in the SE pipeline failed to reveal any systematic differences as a function of temporal distance or in comparison to a non-autobiographical control task, and thus failed to support any specific model in the literature. In contrast, the ME-ICA data supported predictions of the Standard Model of Consolidation regarding both a temporally graded and time-limited role (Squire and Alvarez, 1995; Squire et al., 2015). Moreover, the benefits of ME-ICA were not restricted to the hippocampus. Voxelwise, whole-brain results also demonstrated the improved sensitivity of ME-ICA, both through identification of larger clusters than in the SE data (Figure 2C, Table 2), as well as the addition of several clusters not observed in the SE data. The additional clusters seem unlikely to be spurious findings, but rather captured processes relevant to the experiment: the cluster identified in left frontal cortex has previously been associated with autobiographical recall (with a neurosynth posterior probability of .79), whereas the lateral temporal cluster is typically associated with spoken language (posterior probability = .82).

Naturalistic paradigms, such as the overt cued recall approach described herein, provide promising avenues for researchers to ask questions that are difficult, if not impossible, to address using more limited and controlled laboratory tasks. However, with this flexibility comes additional concerns of how to make the best use of acquired data. Denoising approaches such as ME-ICA may play increasingly important roles in data processing strategies and offer the potential to inform long-standing theoretical debates.

## ACKNOWLEDGEMENTS

This work was supported by the Intramural Research Program at the National Institute of Mental Health (ZIAMH002920) and was conducted under NIH Clinical Study Protocol 93-M-0170 (clinical-trials.gov ID: NCT00001360). The authors thank Andrew Persichetti, Samantha Audrain, and Jason Avery for helpful discussions regarding this work and Bess Bloomer, Sarah Kalinowski, and Alina Quach for assistance with data collection and preparation.

## DATA AVAILABILITY STATEMENT

fMRI data used in this experiment are publicly available at OpenNeuro.org (accession number: ds003511).

## DETAILED METHODS

### Participants

Data for this experiment were taken from a previously published lab dataset (Gilmore et al., 2021a; Gilmore et al., 2021b). Participants consisted of 40 right-handed young adult participants (23 female; mean age = 24.2 years old) who were native English speakers with normal or corrected-to-normal vision and reported no history of psychiatric or neurologic illness. Informed consent was obtained from all participants and the experiment was approved by the NIH Institutional Review Board (clinical trials number NCT00001360). Participants received monetary compensation for their participation.

### Stimuli

Stimuli consisted of 48 photographic images depicting people engaged in various activities. Images were sized at 525 × 395 pixels (screen resolution: 1920 × 1080 pixels) and presented against a black background. Stimuli were presented using PsychoPy2 software (Peirce, 2007; RRID: SCR_006571) on an HP desk-top computer running Windows 10.

### Autobiographical recall task

In this task, participants retrieved and described autobiographical memories in response to photographic picture cues. For each trial, participants were first directed to recall an event from one of 3 different time periods (earlier in the same day [“Today”], 6-18 months prior, or 5-10 years prior). Participants were given a choice of 2 photographic cues and had 11 s to select the picture they preferred, which they indicated via button press response. The screen was replaced with a fixation cross once a response was made, and at the end of the selection period an enlarged version of the selected image was presented in the center of the screen for 5 s. Participants used this period to think back to a specific autobiographical event.

Following picture presentation, participants were given 116 s to describe an event while a white fixation cross was presented centrally. Participants were instructed to describe each event in as much detail as possible for the full duration of each trial. Additional task details are described in (Gilmore et al., 2021a). A 2.2 s red fixation cross indicated the end of each trial, and trials were separated by a 19.8 s fixation period. One trial from each of the 3 time periods was included in each Autobiographical Recall task scan run.

Participants were given practice with the task before scanning, and if the events described were not specific, participants were re-instructed and given further practice until specific episodes were being described. During this time, participants were also instructed not to repeat event descriptions in the experiment.

### Picture description control task

As an active control task, participants described events being depicted in cue photographs. The trial timing and structure was identical to that used for the autobiographical recall task, except participants were instructed during the 5 second picture display period to scrutinize the image so that they could describe it in as much detail as possible when it was removed, rather than use it to recall a memory. As before, trials were separated by 19.8 s of fixation, and three trials were included per picture description control task run.

### Audio recording, transcript scoring, and alignment of spoken responses to the BOLD timeseries

The processing steps associated with the recording and scoring of spoken responses have been described in detail previously (Gilmore et al., 2021a; Gilmore et al., 2021b). Briefly, recorded audio was transcribed and scored for content using an adapted form of the Autobiographical Interview (Levine et al., 2002; Gaesser et al., 2011). This procedure separates “Internal” (episodic) details specific to the event details from other types of “External” details.

Subcategories of Internal details included: Activities, Objects, Perceptual, Person, Place, Thought/Emotion, Time, and Miscellaneous. External detail types included Episodic (i.e., details from other events), Repetitions, Semantic statements, and a catch-all “Other External” category. Timestamps for each spoken word and phrase were generated and matched with the text in transcripts, and different categories of recalled content were converted into event-related regressors for fMRI data analysis, as will be described below.

### fMRI data acquisition

Data were acquired on a General Electric Discovery MR750 3.0T scanner, using a 32-channel phased-array head coil. Functional images were acquired using a BOLD-contrast sensitive multi-echo echo-planar sequence (Array Spatial Sensitivity Encoding Technique [ASSET] acceleration factor = 2, TEs = 12.5, 27.6, and 42.7 ms, TR = 2200 ms, flip angle = 75°, 64 × 64 matrix, in-plane resolution = 3.2 × 3.2 mm). Whole-brain EPI volumes (MR frames) of 33 interleaved, 3.5-mm-thick oblique slices were obtained every 2.2 s. Slices were manually aligned to the AC-PC axis. A high-resolution T1 structural image was also obtained for each subject (TE = 3.47 ms, TR = 2.53 s, TI = 900 ms, flip angle = 7°, 172 slices of 1 × 1 × 1 mm voxels).

Foam pillows were provided for all participants to help stabilize head position and scanner noise was attenuated using foam ear plugs and a noise-cancelling headset. This headset was also used to communicate with the participant during their time in the scanner. Heart rate was recorded via a sensor placed on the left middle finger and a belt monitored respiration.

### fMRI processing: Single echo (standard) analysis

fMRI data were processed following a standard SE preprocessing routine in AFNI (Cox, 1996) (RRID: SCR_005927) to reduce noise and facilitate across-subject comparisons. This processing stream used the 2^nd^ echo, as this TE is within the typical range single echo fMRI paradigms use to study autobiographical recall (e.g., Abraham et al., 2008; Szpunar et al., 2009; Weiler et al., 2010; Gilmore et al., 2018; St. Jacques et al., 2018), including those focusing on the MTL or hippocampus (e.g., Svoboda and Levine, 2009; Bonnici and Maguire, 2012; Thakral et al., 2020). Steps included removal of the first 4 frames to remove potential T1 equilibration effects (*3dTcat*), despiking to remove large transients in the timeseries (*3dDespike*), slice-time correction (*3dTshift*) and frame-by-frame rigid-body realignment to the first volume of each run (*3dvolreg*). Data from each scan run were blurred with a 4mm FWHM smoothing kernel to approximate the smoothness and minimum cluster extents required to maintain a corrected *p* < .05 for whole-brain effects in the ME-ICA pipeline. Data were registered to each individual’s T1 image, normalized by the grand mean of each run, and then resampled into 3-mm isotropic voxels and linearly transformed into Talairach atlas space (Talairach and Tournoux, 1988).

### fMRI processing: Multi-echo ICA

Multi-echo data were also preprocessed using AFNI, using the same procedures described in previous publications of these data (Gilmore et al., 2021a; Gilmore et al., 2021b). Initial steps for each TE of each run were identical to those used in the SE processing stream included a removal of the first 4 frames, despiking, slice-time correction, and rigid-body volume registration. Following these initial steps, data from the three echoes acquired for each run were used to remove additional noise using ME-ICA (Kundu et al., 2012) as implemented in the *meica*.*py* AFNI function.

This procedure initially calculates a weighted average of the different echo times (“optimally combined” data) to reduce thermal noise within each voxel and registers the resulting image to the corresponding anatomical scan. Separately, the echo-specific data are submitted to spatial ICA, and the known properties of T_2_^*^ signal decay over time (across the echoes) are used to separate putatively neural components from artefactual components, such as thermal noise or head motion (Power et al., 2018). To be retained, components must show a strong fit with a model that assumes a temporal dependence on signal intensity and also a poor fit with a model that assumes temporal independence (Kundu et al., 2012). Components determined to be noise are then regressed from the optimally combined data. Selection criteria were left at the default settings of AFNI’s *tedana*.*py* function. Following ME-ICA processing, data were spatially blurred with a Gaussian kernel 3mm full-width at half-maximum, normalized by the grand mean of each run, and then resampled into 3-mm isotropic voxels and linearly transformed into Talairach atlas space (Talairach and Tournoux, 1988), as in the SE pipeline.

### GLM creation

Data from both the SE and ME-ICA streams were linearly detrended and analyzed in AFNI using the same general linear model (GLM) approach (*3dDeconvolve*). The initial picture selection period was modeled using a single HRF across all trial types convolved with a boxcar of 11 s duration. The subsequent Picture Display period was also modeled with a single HRF convolved with a boxcar of 5 s duration. The analysis of recall effects utilized a mixed block/event related design (Visscher et al., 2003). Separate regressors modeled sustained effects related to the narration periods of the Autobiographical Recall and Picture Description narration periods. These convolved an HRF with a boxcar of 118.2 s duration in all cases. Additional regressors coded for transient effects associated with each of the 12 types of detail derived from the Autobiographical Interview scoring procedure as described above. Six motion parameters (3 translational, 3 rotational) were included in each subject’s GLM as regressors of non-interest.

### Comparing SE and ME-ICA effects within the hippocampus

Anterior and posterior regions of the hippocampus were defined for each participant, using the uncal apex as a landmark as described previously (Gilmore et al., 2021a). Activity within each hippocampal ROI was averaged across voxels for each condition, using the Picture Description control task as a baseline.

To determine the effect of the processing pipeline on the observed activity differences related to the temporal distance of each event, a repeated measures ANOVA was constructed. This included factors of temporal distance (2 levels: Today, 5-10 years ago), subregion (2 levels: anterior, posterior), hemisphere (2 levels: left, right), and processing pipeline (2 levels: SE, ME-ICA). Pairwise comparisons were conducted to characterize significant interactions when appropriate.

Activity in each region, for each pipeline, and each temporal distance was compared against the Picture Description baseline task, using one-sample *t*-tests. Due to the large number of comparisons involved, correction for multiple comparisons included a Bonferroni approach as well False Discovery Rate (FDR). The latter was performed in matlab using *fdr_bh*.*m*.

### Voxelwise analysis of temporal distance effects

To test for effects of the recency or remoteness of a memory on retrieval-related BOLD activity, a voxelwise whole brain contrast (paired-samples, two-tailed) of the Today and 5-10 year ago conditions was performed on the SE and ME-ICA pipeline data for each subject.

Correction to a whole brain *p* < .05 was achieved by requiring a voxelwise *p* < .001 and a minimum cluster extent of 17 voxels for the SE data and 18 for the ME-ICA data, determined for each pipeline using AFNI’s *3dClustSim* and its non-Gaussian (-*acf*) autocorrelation function (Cox et al., 2017). Both pipelines identified large (<800 voxel) posterior midline clusters containing 3 distinct local maxima. Center of mass coordinates were identified for each location by incrementing the voxelwise *t*-statistic threshold in steps of 0.1 until the 3 clusters were separated. This was achieved at *t* > 5.04 for the SE data and *t* > 5.36 for the ME-ICA data.

## REFERENCES

Abraham A, Schubotz RI, Von Cramon DY. (2008). Thinking about the future versus the past in personal and non-personal contexts. Brain Research 1233:106–119. 10.1016/j.brainres.2008.07.084.

Alvarez P, Squire LR. (1994). Memory consolidation and the medial temporal lobe: A simple network model. Proc Natl Acad Sci USA 91:7041–7045.

Barry DN, Maguire EA. (2019). Remote memory and the hippocampus: A constructive critique. Trends Cog Sci 23:128–142. 10.1016/j.tics.2018.11.005.

Benjamini Y, Yekutieli D. (2001). The control of the false discovery rate in multiple testing under dependency. Annals of Statistics 29:1165–1188. 10.1214/aos/1013699998.

Bonnici HM, Maguire EA. (2012). Detecting representations of recent and remote autobiographical memories in vmPFC and hippocampus. J Neurosci 32:16982–16991. 10.1523/jneurosci.2475-12.2012.

Caballero-Gaudes C, Reynolds RC. (2017). Methods for cleaning the BOLD fMRI signal. Neuroimage 154:128–149. 10.1016/j.neuroimage.2016.12.018.

Chen J, Leon gYC, Honey CJ, Yong CH, Norman KA, Hasson U. (2017). Shared memories reveal shared structure in neural activity across individuals. Nat Neurosci 20:115–125. 10.1038/nn4450.

Cox RW. (1996). AFNI: software for analysis and visualization of functional magnetic resonance images. Comput Biomed Res 29:162–173. 10.1006/cbmr.1996.0014.

Cox RW, Chen G, Glen DR, Reynolds RC, Taylor PA. (2017). fMRI clustering in AFNI: False-positive rates redux. Brain Connect 7:152–171. 10.1089/brain.2016.0475.

Finn ES, Corlett PR, Chen G, Bandettini PA, Constable RT. (2018). Trait paranoia shapes inter-subject synchrony in brain activity during an ambiguous social narrative. Nature Communications 9:2043. 10.1038/s41467-018-04387-2.

Gaesser B, Sacchetti DC, Addis DR, Schacter DL. (2011). Characterizing age-related changes in remembering the past and imagining the future. Psychol Aging 26:80. 10.1037/a0021054.

Gilboa A, Moscovitch M. (2021). No consolidation without representation: Correspondence between neural and psychological representations in recent and remote memory. Neuron 109:2239–2255. 10.1016/j.neuron.2021.04.025.

Gilmore AW, Nelson SM, Chen H-Y, McDermott KB. (2018). Task-related and resting-state fMRI identify distinct networks that preferentially support remembering the past and imagining the future. Neuropsychologia 110:180–189. 10.1016/j.neuropsychologia.2017.06.016.

Gilmore AW, Quach A, Kalinowski SE, González-Araya EI, Gotts SJ, Schacter DL, Martin A. (2021a). Evidence supporting a time-limited hippocampal role in retrieving autobiographical memories. Proc Natl Acad Sci USA 118:e2023069118. 10.1073/pnas.2023069118.

Gilmore AW, Quach A, Kalinowski SE, Gotts SJ, Schacter DL, Martin A. (2021b). Dynamic content reactivation supports naturalistic autobiographical recall in humans. J Neurosci 41:153–166. 10.1523/jneurosci.1490-20.2020.

Gonzalez-Castillo J, Panwar P, Buchanan LC, Caballero-Gaudes C, Handwerker DA, Jangraw DC, Zachariou V, Inati S, Roopchansingh V, Derbyshire JA, Bandettini PA. (2016). Evaluation of multi-echo ICA denoising for task based fMRI studies: Block designs, rapid event-related designs, and cardiac-gated fMRI. Neuroimage 141:452–468. 10.1016/j.neuroimage.2016.07.049.

Gracco VL, Tremblay P, Pike B. (2005). Imaging speech production using fMRI. Neuroimage 26:294–301. 10.1016/j.neuroimage.2005.01.033.

Hasson U, Honey CJ. (2012). Future trends in neuroimaging: Neural processes as expressed within real-life contexts. Neuroimage 62:1272–1278. 10.1016/j.neuroimage.2012.02.004.

Haxby JV, Gobbini MI, Nastase SA. (2020). Naturalistic stimuli reveal a dominant role for agentic action in visual representation. Neuroimage 216:116561. 10.1016/j.neuroimage.2020.116561.

Jasmin K, Gotts SJ, Xu Y, Liu S, Riddell CD, Ingeholm JE, Kenworthy L, Wallace GL, Braun AR, Martin A. (2019). Overt social interaction and resting state in young adult males with autism: Core and contextual neural features. Brain 142:808–822. 10.1093/brain/awz003.

Kundu P, Brenowitz ND, Voon V, Worbe Y, Vértes PE, Inati SJ, Saad ZS, Bandettini PA, Bullmore ET. (2013). Integrated strategy for improving functional connectivity mapping using multiecho fMRI. Proc Natl Acad Sci USA 110:16187–16192. 10.1073/pnas.1301725110.

Kundu P, Inati SJ, Evans JW, Luh W-M, Bandettini PA. (2012). Differentiating BOLD and non-BOLD signals in fMRI time series using multi-echo EPI. Neuroimage 60:1759–1770. 10.1016/j.neuroimage.2011.12.028.

Kundu P, Voon V, Balchandani P, Lombardo MV, Poser BA, Bandettini PA. (2017). Multi-echo fMRI: A review of applications in fMRI denoising and analysis of BOLD signals. Neuroimage 154:59–80. 10.1016/j.neuroimage.2017.03.033.

Levine B, Svoboda E, Hay JF, Winocur G, Moscovitch M. (2002). Aging and autobiographical memory: Dissociating episodic from semantic retrieval. Psychol Aging 17:677-689. 0.1037//0882-7974.17.4.677.

Nadel L, Moscovitch M. (1997). Memory consolidation, retrograde amnesia and the hippocampal complex. Curr Opin Neurobiol 7:217–227. 10.1016/S0959-4388(97)80010-4.

Power JD, Lynch CJ, Gilmore AW, Gotts SJ, Martin A. (2019). Reply to Spreng et al.: Multiecho fMRI denoising does not remove global motion-associated respiratory signals. Proc Natl Acad Sci USA 116:19243–19244. 10.1073/pnas.1909852116.

Power JD, Plitt M, Gotts SJ, Kundu P, Voon V, Bandettini PA, Martin A. (2018). Ridding fMRI data of motion-related influences: Removal of signals withdistinct spatial and physical bases in multiecho data. Proc Natl Acad Sci USA 115:E2105–E2114. 10.1073/pnas.1720985115.

Sekeres MJ, Winocur G, Moscovitch M. (2018). The hippocampus and related neocortical structures in memory transformation. Neurosci Lett 680:39–53. 10.1016/j.neulet.2018.05.006.

Siegel JS, Power JD, Dubis JW, Vogel AC, Church JA, Schlaggar BL, Petersen SE. (2013). Statistical improvements in functional magnetic resonance imaging analyses produced by censoring high-motion data points. Hum Brain Mapp 35:1981–1996. 10.1002/hbm.22307.

Spreng RN, Fernánzez-Cabello S, Turner GR, Stevens WD. (2019). Take a deep breath: Multiecho fMRI denoising effectively removes head motion artifacts, obviating the need for global signal regression. Proc Natl Acad Sci USA 116:19241–19242. 10.1073/pnas.1909848116.

Squire LR, Alvarez P. (1995). Retrograde amnesia and memory consolidation: a neurobiological perspective. Curr Opin Neurobiol 5:169–177.

Squire LR, Genzel L, Wixted JT, Morris RG. (2015). Memory consolidation. Cold Spring Harb Perspect Biol 7:a021766. 10.1101/cshperspect.a021766.

St. Jacques Pl, Carpenter AC, Szpunar KK, Schacter DL. (2018). Remembering and imagining alternative versions of the personal past. Neuropsychologia 110:170–179. 10.1016/j.neuropsychologia.2017.06.015.

Svoboda E, Levine B. (2009). The effects of rehearsal on the functional neuroanatomy of episodic autobiographical and semantic remembering: A functional magnetic resonance imaging study. J Neurosci 29:3073–3082. 10.1523/JNEUROSCI.3452-08.2009.

Szpunar KK, Chan JCK, McDermott KB. (2009). Contextual processing in episodic future thought. Cereb Cortex 19:1539–1548. 10.1093/cercor/bhn191.

Talairach J, Tournoux P. (1988). Co-planar stereotaxic atlas of the human brain. New York: Thieme Medical Publishers, Inc.

Thakral PP, Madore KP, Addis DR, Schacter DL. (2020). Reinstatement of event details during episodic simulation in the hippocampus. Cereb Cortex 30:2321–2337. 10.1093/cercor/bhz244.

Visscher KM, Miezin FM, Kelly JE, Buckner RL, Donaldson DI, McAvoy MP, Bhalodia VM, Petersen SE. (2003). Mixed blocked/event-related designs separate transient and sustained activity in fMRI. Neuroimage 19:1694–1708. 10.1016/S1053-8119(03)00178-2.

Weiler J, Suchan B, Daum I. (2010). When the future becomes the past: Differences in brain activation patterns for episodic memory and episodic future thinking. Behav Brain Res 212:196–212.

Yonelinas AP, Ranganath C, Ekstrom AD, Wiltgen BJ. (2019). A contexual binding theory of episodic memory: systems consolidation reconsidered. Nat Rev Neurosci 20:364–375. 10.1038/s41583-019-0150-4.

